# Synchronization of sensory gamma oscillations promotes multisensory communication

**DOI:** 10.1101/523688

**Authors:** Jonas Misselhorn, Bettina C. Schwab, Till R. Schneider, Andreas K. Engel

## Abstract

Rhythmic neuronal activity in the gamma range is a signature of active cortical processing and its synchronization across distant sites has been proposed as a fundamental mechanism of network communication. While this has been shown within sensory modalities, we tested whether crosstalk between the senses relies on similar mechanisms. In two consecutive experiments, we used a task in which human participants (male and female) matched amplitude changes of concurrent visual, auditory and tactile stimuli. In this task, matching of congruent stimuli was associated with a behavioral benefit compared to matching of incongruent stimuli. In the first experiment, we used source-level analysis of high-density electroencephalography (EEG) and observed that cross-modal matching of congruent inputs was associated with relatively weaker gamma band coherence between early sensory regions. Next, we used bifocal high-definition transcranial alternating current stimulation (hd-tACS) to manipulate the strength of coupling between sensory cortices. Here, we used a lateralized version of the task in which hd-tACS was applied either ipsilateral or contralateral to the hemisphere receiving sensory stimuli. Ipsilateral gamma, but not alpha stimulation slowed responses to congruent trials whereas responding to incongruent trials was not changed by hd-tACS. We speculate that fast responding to congruent stimuli involves decoupling of sensory gamma oscillations, which was prevented by hd-tACS. Collectively, these results indicate that coordinated sensory gamma oscillations play an important role for direct cross-modal interactions. We suggest that, comparable to interactions within sensory streams, phase-coupled gamma oscillations might provide the functional scaffold for cross-modal communication.

**Significance statement:** Cortical gamma oscillations structure segregated neural activity and were suggested to represent a fundamental mechanism of network communication. While there is ample evidence for the role of long-range gamma synchronization in unisensory processing, its significance in multisensory networks is still unclear. We show that coordinated sensory gamma oscillations play an important role for direct cross-modal interactions and propose that phase synchronization promotes communication between sensory cortices. To that end, we carried out two consecutive experiments using state-of-the-art high-density electroencephalography (EEG) and high-definition transcranial alternating current stimulation (hd-tACS). By complementing an observational with an interventional method, we provide novel evidence for the role of synchronized gamma oscillations in multisensory communication.

## Introduction

Perceiving the world through distinct sensory channels provides complementary as well as redundant and conflicting information about the environment. In order to structure these sensory signals, fundamental neuronal computations are concerned with cross-modal matching of sensory signals. On the neuronal and behavioral level, processing cross-modally congruent stimuli is associated with enhanced efficiency when compared with incongruent or unimodal processing and often coincides with enhanced cortical activity (Ghanzafar and Schroeder, 2006) and behavioral benefits (Spence, 2011). Within sensory systems, such integrative processes likely involve corticocortical synchronization of high-frequency oscillatory activity (Engel et al., 2001; Fries, 2009). For instance, perceptual grouping and feature binding across cortical columns and hemispheric homologues of visual cortex have been shown to involve phase coupling of neuronal gamma band oscillations (Gray et al., 1989; Engel et al., 1991). Relatedly, it was suggested that synchronized oscillations might provide a solution to the binding problem (Tallon-Baudry et al., 1996; Treisman, 1996). Moreover, gamma oscillations have been proposed to constitute a framework that allows carrying cohesive patterns of neural activity along sensory streams (Fries, 2015). Taken together, the coordination of gamma oscillations may enable structuring as well as transmitting sensory information within sensory networks and thereby likely plays an important role in orchestrating multisensory interactions.

A number of studies have investigated gamma band activity during multisensory perception. Visual stimulus detection, for instance, was shown to be improved by redundant auditory stimuli while gamma band responses in frontal cortex were enhanced (Senkowski et al, 2005, 2007). Recognition and classification of visual objects was improved by congruent auditory input showing increased gamma band power in temporal or parietal cortices (Yuval-Greenberg et al. 2007; Schneider et al., 2008a). While the aforementioned studies showed multisensory modulations of gamma band power in association cortices, other studies also noted changes in sensory cortices (Krebber et al., 2015; Friese et al., 2016). For instance, attention for supra-threshold audio-visual stimuli was associated with enhanced sensory gamma oscillations in both visual and auditory cortex (Friese et al., 2016) and matching congruent visual-tactile motion stimuli induced enhanced gamma power in visual and somatosensory cortex (Krebber et al., 2015). Additionally, there is evidence for altered gamma oscillations underlying schizophrenia (Uhlhaas and Singer, 2010; Curic et al., 2019). In these patients, aberrant multisensory integration was shown to be accompanied by altered gamma band dynamics in response to multisensory stimuli (Stone et al., 2014; Balz et al. 2016). Taken together, cross-modally corresponding or congruent stimuli typically induce strong local synchronization of gamma band oscillations in both sensory and associations cortices.

In addition to local changes of gamma band activity, it was suggested that cross-modal interactions involve inter-areal phase synchronization of sensory gamma oscillations (Senkowski et al., 2008). Specifically, enhanced processing of cross-modally congruent stimuli might imply feature binding across modalities mediated by synchronization of sensory gamma oscillations. A constraint in testing this hypothesis is that differences in power constitute a bias for the computation of phase coherence (Bastos and Schoffelen, 2016). As reviewed above, many multisensory paradigms would thus not be suited for testing this prediction. Here, we present two consecutive experiments designed to (1) observe patterns of functional connectivity between sensory cortices during cross-modal matching in human electroencephalogram (EEG) and (2) test the behavioral relevance of this functional coupling with bifocal high-definition transcranial alternating current stimulation (hd-tACS). First, we re-analyzed EEG data recorded during a cross-modal matching paradigm where no gamma power changes were observed (Anonymous, 2018). In this task, participants matched concurrent amplitude changes of visual, auditory and tactile stimuli that were either congruent (increases or decreases in stimulus intensity of two modalities) or incongruent (increase in one and decrease in the other modality). We hypothesized the behavioral benefit of cross-modal congruence would be accompanied by increased sensory gamma coupling. Surprisingly, we found the opposite pattern of results, i.e., enhanced gamma band coherence for incongruent stimuli. In a second study, we tested the functional relevance of this finding by means of bifocal hd-tACS designed to target coupling between visual and somatosensory cortex in a visual-tactile version of the amplitude matching task. Electrical stimulation was applied at either 10 or 40 Hz with 0° or 180° phase shift between montages. Also, both sensory and electrical stimulation were applied lateralized resulting in ipsilateral stimulation (tACS was applied over cortices targeted by sensory stimuli) and contralateral stimulation regimes. We hypothesized that only ipsilateral stimulation would induce task-specific effects while contralateral stimulation served as an active control. Specifically, we expected a reduction in congruence-related behavioral benefits under ipsilateral in-phase 40 Hz stimulation.

## Methods

### EEG experiment

#### Participants

Twenty-one participants (11 female, 23.8 ± 2.5 years) were invited for two sessions of EEG. None of them had a history of neurological or psychiatric disorders and visual, auditory and tactile perception were normal or corrected to normal. The experiment was approved by the ethics committee of the Hamburg Medical Association, all participants gave written consent and received monetary compensation for their participation.

#### Experimental design

Participants received trimodal sensory stimulation (see *Stimulus material* for details) on each trial of the experiment. These trimodal stimuli contained a visual, an auditory and a tactile component. On each trial, all components underwent a brief intensity change; that is, visual contrast, auditory loudness and vibration strength were either increased or decreased. The task was to attend bimodal pairs (VT, visual-tactile or AV, audio-visual) blockwise and compare attended intensity changes (Fig. 1A). These changes could be either congruent (i.e., in the same direction) or incongruent (i.e., in different directions), the respective third modality had to be ignored. Participants responded verbally after stimulus offset. Due to the forced wait period, response times could not be analyzed. Blocks of VT and AV attention contained 64 trials with equal contributions of the eight possible stimulus configurations of increases and decreases across modalities. On two separate days, 10 blocks of each VT and AV attention were performed in an alternating fashion summing up to 1280 trials.

**Figure 1.**
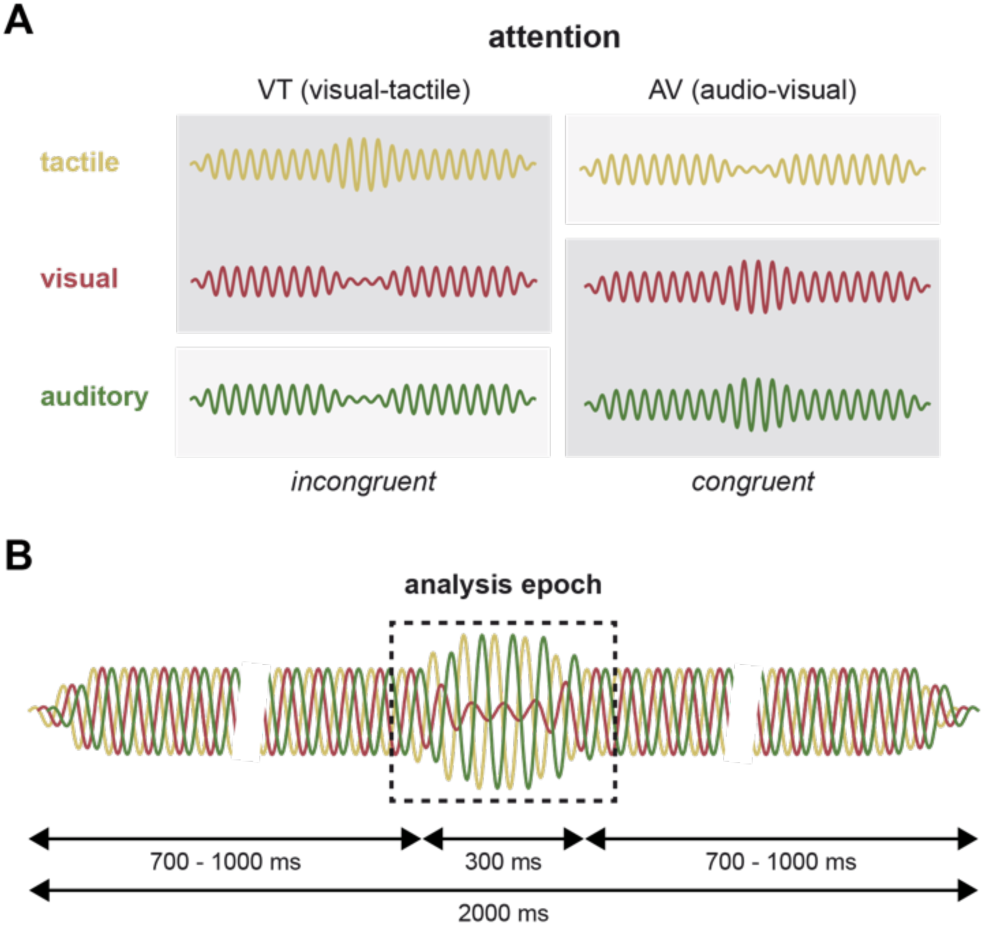
Experimental design of EEG experiment. **(A)** In different blocks, participants attended visual-tactile (VT) or audio-visual (AV) pairs out of a trimodal stimulus. Their task was to report whether the attended stimulus components changed congruently or incongruently. **(B)** Trimodal stimuli had a fixed duration of 2000 ms with a jittered onset of the amplitude change which lasted for 300 ms. Analyses were performed for the period of change.

#### Stimulus material

Visual stimulation consisted of a circular, expanding grating presented centrally on a CRT screen (Iiyama, Model HM204DTA, refresh rate: 120 Hz) with grey background at a visual angle of 5°. The auditory stimulus component was a complex sinusoidal tone (13 sine waves: 64 Hz and its first 6 harmonics as well as 91 Hz and its first 5 harmonics, low-frequency modulator: 0.8 Hz) played back with audiometric insert earphones binaurally at 70 dB (E-A-RTONE 3A, 3M, USA). The tactile component was a high-frequency vibration delivered to the fingertips of both index fingers (250 Hz on C2 tactors, Engineering Acoustics Inc., USA). Visual contrast, auditory loudness and vibration amplitude were experimentally modulated. In total, trimodal stimuli had a fixed duration of 2 s and changes in intensity lasted for 300 ms (Fig. 1B). Transitions were smoothed with cosine tapers and onsets were jittered across trials between 700 and 1000 ms after stimulus onset. The magnitude of change per modality and change direction was estimated individually with a psychometric step-function prior to experimental blocks on each day (Watson and Pelli, 1983).

#### EEG recordings

High-density EEG was recorded from 128 channels using active Ag/AgCl electrodes referenced to the nose (EasyCap, Germany) via BRAINAMP MR amplifiers (Brain Products, Germany) and digitized after analog filtering (low cutoff: 10 s, high cutoff: 1000 Hz, sampling rate: 1000 Hz). After down-sampling to 500 Hz, data was filtered (high-pass: 1 Hz, low-pass: 120 Hz, notch: 49-51 Hz, 99-101 Hz) and cut into epochs locked to stimulus onset (−500 to 2000 ms). Prior to pre-processing, EEG was re-referenced to the common average. Dual-band independent components analysis (ICA) was used to remove stereotypical artifacts including eye blinks, saccades, electrocardiogram and other myogenic activity (lower band: 1-30 Hz, higher band: 30-120 Hz). Due to low signal-to-noise ratio, 19 electrodes of the outer rim covering neck and chin were removed. Stratified data held on average 426 ± 89 epochs per participant. In sensor space, event-related potentials were averaged per experimental condition and subtracted from single-trial data. Source reconstruction was performed with exact low-resolution electromagnetic tomography (eLORETA, regularization: 0.05; Pascual-Marqui et al. 2011). Spatial filters were constructed using a three-shell head model (Nolte and Dassios, 2005) and a cortical grid in MNI space obtained by down-sampling the Freesurfer template to 10000 grid points (Desikan et al., 2006). Dipole directions were separately for the chosen frequencies of interest (see *Data analysis*) by maximizing spectral power with singular value decomposition.

#### Data analysis

Frequencies of interest in alpha/beta and gamma bands were defined by computing the global interaction measure (GIM) in sensor space (Ewald et al., 2012). GIM quantifies the overall strength of connectivity across all connections and yields a full spectrum, allowing to identify frequencies of maximal coupling. We computed GIM with a frequency resolution of 1 Hz based on whole trial data from all conditions (Fig. 2-1). We defined peak frequencies for alpha/beta and gamma bands as the maximum between 8 and 20 Hz (alpha/beta) respectively 60 and 90 Hz (gamma). Individual peak frequencies for alpha/beta ranged from 9 to 16 Hz (13.19 ± 1.66 Hz) and for gamma from 60 to 90 Hz (77.19 ± 9.63 Hz). In source space, cross-spectra between all cortical grid points were computed at the frequencies of interest identified by GIM in a time window of 500 ms centered on the change. Imaginary coherence (iCoh) and power were computed by fast fourier transform using Hanning windows (Nolte et al., 2004). Regions of interest (ROI) for primary visual, auditory and somatosensory cortex were defined anatomically by reference to the Freesurfer atlas (Desikan et al., 2006) and iCoh was averaged in bimodal networks for all edges between respective ROIs (i.e., visual-tactile, audio-visual and audio-tactile). In order to analyze systematic biases to the computation of iCoh, we estimated local synchronization of cortical activity in the alpha/beta and gamma band. Event-related power was averaged for each ROI and subjected to analysis of variance (ANOVA) with factors ROI (visual/auditory/somatosensory), *ATTENTION* (VT/AV) and *CONGRUENCE* (congruent/incongruent) separately for alpha/beta and gamma bands. Similarly, iCoh was analyzed by ANOVA with factors *NETWORK* (visual-tactile/audio-visual/audio-tactile), *ATTENTION* (VT/AV) and *CONGRUENCE* (congruent/incongruent) separately for alpha/beta and gamma bands. Where necessary, Greenhouse-Geisser correction was applied. Tables containing complete results from ANOVA is provided as extended data (Fig. 2-2).

**Figure 2.**
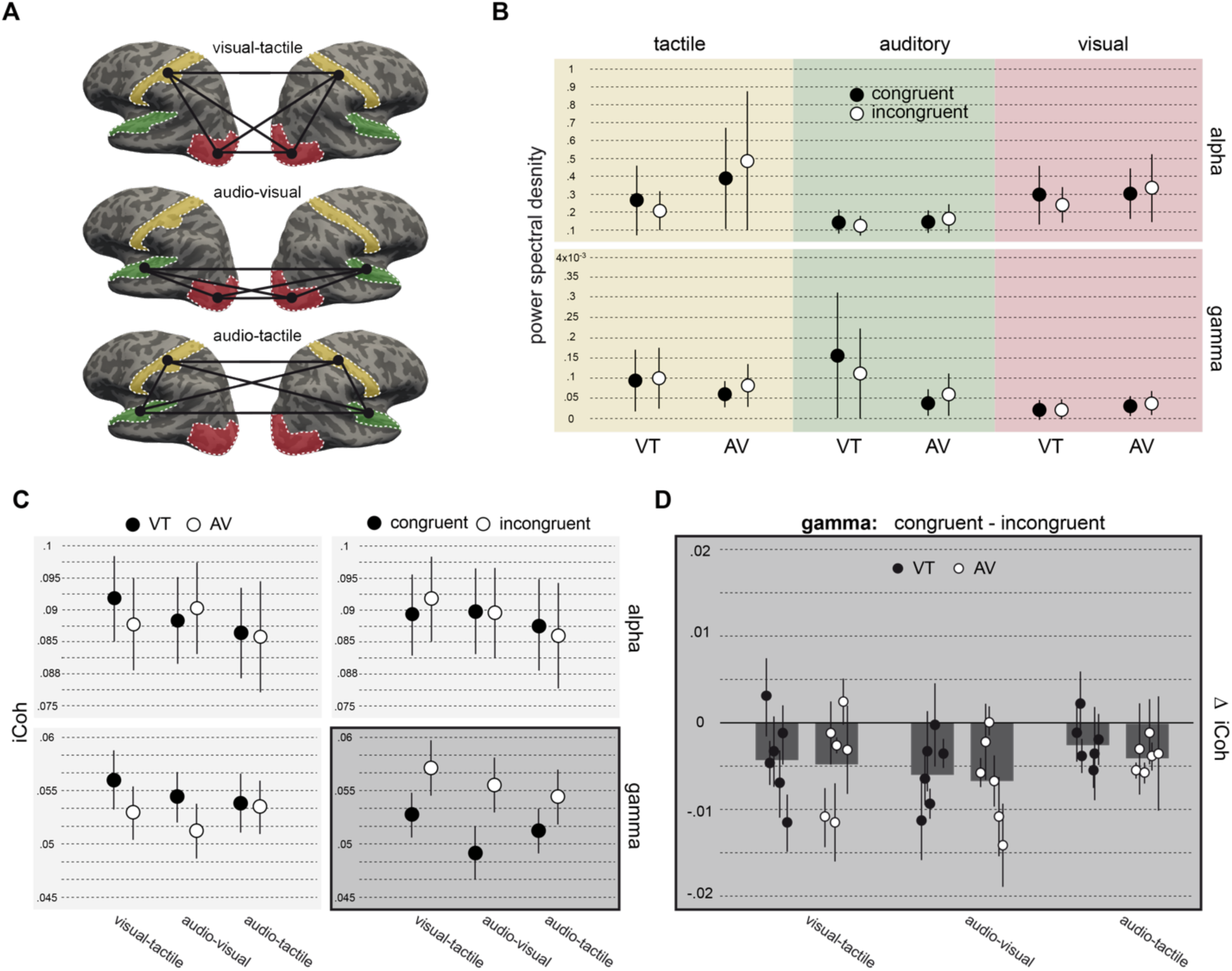
Results EEG experiment. **(A)** Schematic of analyzed network edges. Regions of interest (ROI) were based on the Freesurfer atlas (Red = visual cortex, green = auditory cortex, yellow = somatosensory cortex). **(B)** No effects of *ATTENTION* or *CONGRUENCE* on power in the alpha/beta and gamma bands for each ROI (same color coding as in (A)). Individual frequencies for the two bands were determined by global interaction measure (see *Methods* and Supplement 1). Black circles = congruent presentations, white circles = incongruent. **(C)** Absolute imaginary coherence (iCoh), averaged over all edges of each network, shows a significant effect of *CONGRUENCE* in the gamma band only. **(D)** Effect of congruence displayed for individual edges of each network and attentional conditions.

### tACS experiment

#### Participants

Twenty-four participants, who had not been enrolled in the EEG experiment, were recruited. All completed a training session after which four participants dropped out due to insufficient performance (<60 % accuracy). Twenty participants completed all experimental sessions (13 female, 25.3 ± 4.5 years). None of them had a history of neurological or psychiatric disorders and visual, auditory and tactile perception were normal or corrected to normal. All gave written consent and received monetary compensation after completion of all three sessions. The Hamburg Medical Association approved the experiment.

#### Experimental design

The task of the tACS experiment was a simplified version of the task employed in the EEG experiment. Details of stimulus material outlined above apply here as well. Instead of trimodal stimuli, only visual-tactile stimulus pairs were presented in the right or left hemifield and to the right or left hand, respectively. Attention was cued by a centrally presented arrow prior to each trial and hemifields were chosen randomly but evenly throughout the experimental blocks. At all times, participants maintained fixation on a central fixation cross. Again, participants were asked to evaluate congruency of change directions. After a training session, participants completed two identical experimental sessions containing three blocks holding 192 trials. Experimental session used either alpha (10 Hz) or gamma (40 Hz) stimulation. These canonical stimulation frequencies were chosen because previous studies showed behavioral as well as neurophysiological effects for these frequencies (Helfrich et al., 2014; Schwab et al., 2018). The order of experimental sessions was counterbalanced across participants. Experimental blocks featured in-phase, anti-phase or sham stimulation (see *Electrical stimulation* for details). The order of stimulation conditions was counter-balanced across participants.

#### Electrical stimulation

Alternating currents were administered in 4-in-1 montages with current flow between four outer and one central electrode (Patel et al., 2009; Saturnino et al., 2015) using Ag/AgCl ring electrodes (diameter = 12 mm). This configuration results in focal electric fields with peaks underneath the central electrode (Fig. 3A). For each participant, we prepared two of these montages designed to target primary visual and primary somatosensory cortex of one hemisphere, respectively. The side of stimulation was counterbalanced across participants. In conjunction with the lateralized experimental design, this resulted in equal proportions of trials that have electrical stimulation contra-or ipsilateral to the hemisphere receiving sensory stimulation (Fig. 3B). Prior to experimental blocks, stimulation was ramped up to 2mA peak-to-peak within 10 s. Sham blocks started with the same ramps, but included no stimulation thereafter. For in-phase stimulation, we used the same waveforms for both montages. For anti-phase stimulation, one waveform was shifted by 180°. In order to prevent inter-montage currents, two separate DC-stimulators were used (DC-Stimulator Plus, Neuroconn, Germany). Stimulators were operated in external mode allowing to control current output via voltage input. The voltage signal was computed in Matlab and produced by a NI-DAQ device run with Labview (NI USB 6343, National Instruments, USA). Impedances of each of the four outer electrodes relative to the central electrode were kept comparable within montages (10-100 kΩ). This is crucial because identical impedances were assumed for the simulation of electric fields.

**Figure 3.**
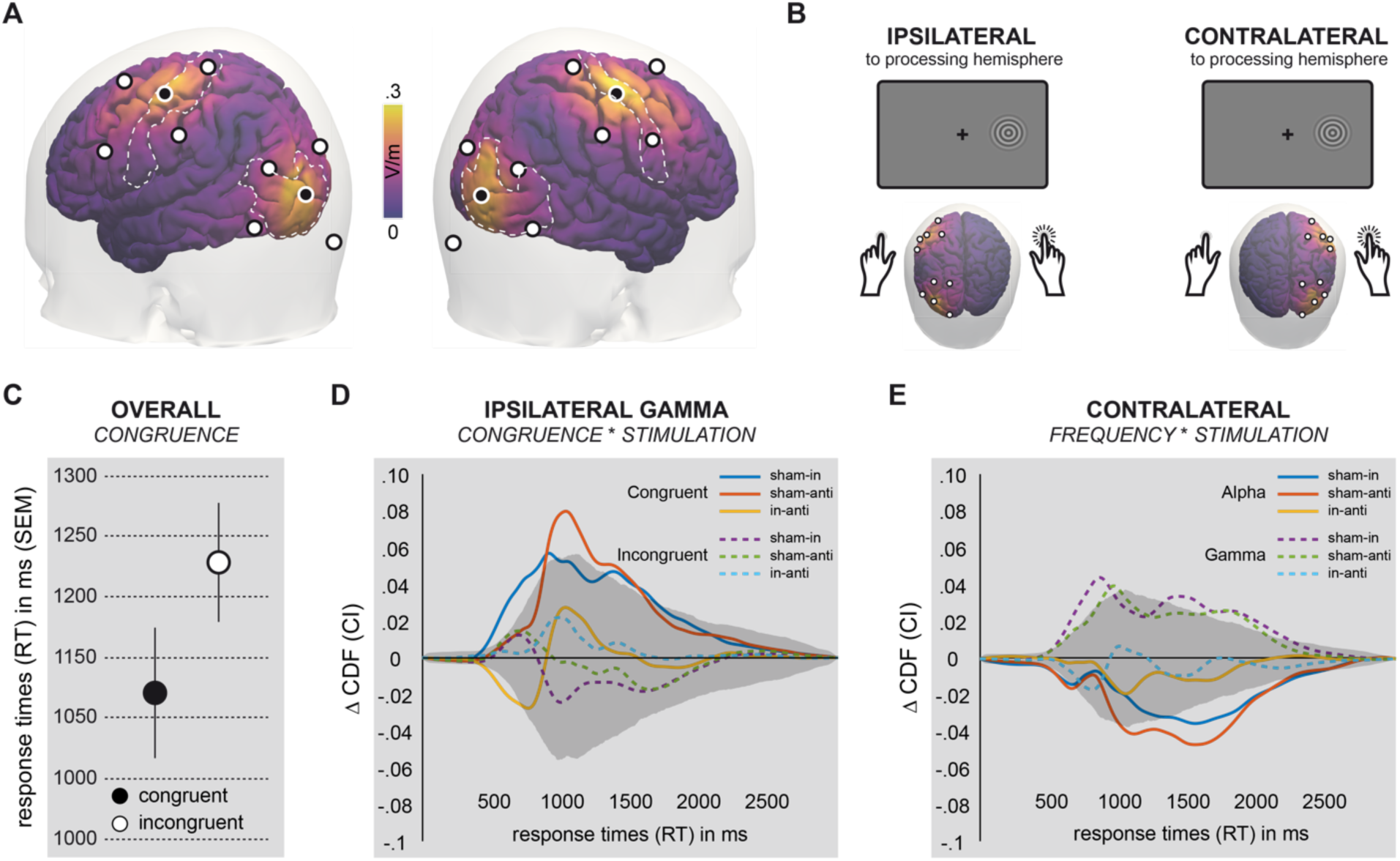
Results tACS experiment. **(A)** Electric fields distributions of bifocal hd-tACS over visual and somatosensory cortex. Alternating currents were applied with two separate left or right hemispheric 4-in-1 montages. Color coding on cortical surface corresponds to the simulated absolute field strength in V/m**. (B)** Schematic showing that visual-tactile and electrical stimulation could be either ipsilateral (hypothesized to be effective with respect to the task) or contralateral (active control) with respect to each other. **(C)** Behavioral effect of *CONGRUENCE* (error bars denote standard error of the mean). **(D)** Significant interaction effect between *CONGRUENCE* (congruent/incongruent) and *STIMULATION* (sham/in-phase/anti-phase) for ipsilateral gamma stimulation was followed up by comparisons of respective cumulative distribution functions (CDFs). Lines represent differences between conditions. Shaded region corresponds to the confidence interval constructed by permutation statistics (alpha = .000129, see *Methods* for details). **(E)** Significant interaction effect between *FREQUENCY* (congruent/incongruent) and *STIMULATION* (sham/in-phase/anti-phase) for contralateral stimulation was followed up by comparisons of respective CDFs. Lines represent differences between conditions. Shaded region corresponds to the confidence interval constructed by permutation statistics (alpha = .000129, see *Methods* for details).

#### Simulation of electric fields

Electrode positions for the 4-in-1 montages were chosen such that electric field strength was maximized in visual and somatosensory areas. For the simulations, we used the same standard cortical MNI model and leadfield matrix *L* that had been used for the source-level analysis of EEG data. We then calculated the electric field 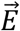 at location 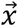 by linear weighting of the leadfield matrix with the injected currents *α*_*i*_, where i denotes indices of the ten stimulation electrodes:

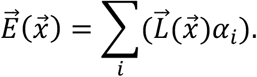

Within visual and somatosensory regions, peak values of 0.3 V/m were reached using currents with peak values of 1 mA. Spatial specificity was high as field strengths rapidly decreased when moving away from the central electrode. This ensured that electrical stimulation was confined to the targeted regions of one hemisphere only.

#### Data analysis

The effects of tACS were evaluated by analyzing response times (RTs). First, we computed an ANOVA with factors *HEMISPHERE* (ipsilateral/contralateral), *FREQUENCY* (alpha/gamma), *STIMULATION* (sham/in-phase/anti-phase) and *CONGRUENCE* (congruent/incongruent). Interactions were followed up by computing reduced ANOVA models. Where necessary, Greenhouse-Geisser correction was applied. Tables containing complete results from ANOVA are provided as extended data (Fig. 3-1). Remaining interactions were followed up by a non-parametric post-hoc analysis. To that end, we estimated cumulative density functions (CDFs) of RT distributions using a Gaussian kernel estimator (Botev et al., 2010). CDFs were estimated for RTs between 0 and 4 s using 1024 bins for each sub-condition and participant. Next, we computed relevant differences between CDFs and averaged across participants. In order to decide about statistical significance of differences in CDFs, we constructed confidence intervals (CIs) by permutation tests. That is, we shuffled all data from a given interaction into two sets, computed CDFs and stored the difference between the CDFs of the two sets as the null-distribution (100.000 permutations). CIs were determined by finding percentiles at each RT bin of the CDF range (lower bound: alpha/2, upper bound: 100-alpha/2). Alpha-level was set to 5% and Bonferroni-corrected for multiple comparisons. Additionally, we corrected for multiple comparisons with respect to the number of RT bins used for CDF. The goal was to expand the CI until only alpha % of all computed differences in the null distribution would fall outside of the CI at any RT bin.

#### Questionnaire of side effects

After each stimulation condition, participants completed a questionnaire designed to reflect (1) the perceived maximum intensity of skin sensations (itching, warmth, stinging, pulsating), phosphenes, fatigue and pain (ranked as either “absent”/0, “light”/1, “moderate”/2, “pronounced”/3 or “strong”/4) as well as (2) the time-course of sensations (“beginning”, “end”, “always”). Condition differences in perceived intensity were evaluated using Wilcoxon matched-pairs signed-rank tests without applying a correction for multiple comparisons in order to maximize power for detecting possibly biasing differences between conditions. Skin sensations were aggregated by computing median responses. In order to analyze whether participants were in fact blinded or whether they could perceive the difference between sham and verum, we computed a binary score reflecting whether participants perceived peripheral sensations only in the beginning (0) or all the time (1). We report averages that can be interpreted as fractions and uncorrected p-values from McNemar tests. Finally, we assessed whether significant tACS-related behavioral effects detected in the main analysis could be explained by the perceived intensity of sensations. To that end, we ranked individual behavioral effects and correlated these scores with the questionnaire data by means of Spearman correlations.

## Results

### Local and long-range synchronization of sensory oscillations

We analyzed power in and coherence between alpha/beta and gamma oscillations in cortical sensory areas. Source reconstructions of scalp EEG were analyzed in an epoch containing the whole change period (Fig. 1B). In order to capture sensory processes with reasonable sensitivity, we selected large ROIs for visual, auditory and somatosensory cortex based on the anatomical Freesurfer parcellation (Desikan et al., 2006). First, we computed raw power for alpha/beta and gamma bands within these regions to exclude biases for the quantification of functional coupling by absolute imaginary coherence (iCoh). Results of ANOVA with factors *ATTENTION* (VT/AV) and *CONGRUENCE* (congruent/incongruent) showed that power in both alpha/beta and gamma bands was comparable across ROIs and experimental conditions (all *p* > .05; Fig. 2B, Fig. 2-1). Second, we computed iCoh between all ROIs and subsequently averaged all edges of visual-tactile, audio-visual and audio-tactile networks (Fig. 2A). Based on the averaged networks, we computed an ANOVA on iCoh values. We found that gamma band coherence differed significantly by factor *CONGRUENCE* only (F(1,20) = 6.666, *p* = 0.018, 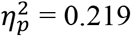; Fig. 2-2), revealing that synchronization in all networks was reduced for congruent relative to incongruent trials (Fig. 2C,D). Although non-significant, larger absolute differences were observed for the task-relevant audio-visual and visual-tactile networks.

### Modulation of behavior with bifocal hd-tACS

In order to modulate functional connectivity between sensory cortices, we applied in-or anti-phase tACS between visual and somatosensory cortex (0°/180° phase shift; Fig. 3A) with either 10 or 40 Hz (2 mA peak-to-peak). Applying electrical fields unilaterally while visual-tactile stimulation was presented lateralized, hd-tACS could be applied either ipsilaterally or contralaterally with respect to the hemisphere receiving sensory stimuli (Fig. 3C,D). The behavioral effects of stimulation were analyzed with ANOVA comprising factors *HEMISPHERE* (ipsilateral/contralateral), *FREQUENCY* (alpha/gamma), *STIMULATION* (sham/in-phase/anti-phase) and *CONGRUENCE* (congruent/incongruent). Participants were well trained on the task and gave, on average, correct responses in ~83 % of all trials. Accuracy differed significantly between congruent and incongruent trials (F(1,19) = 10.122, p = 0.005, 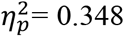; Fig. 3-1), errors occurred less likely in congruent trials (85.48 %) when compared with incongruent trials (80.36 %). Timing of responses showed a similar, but stronger effect of *CONGRUENCE* (Fig. 3C; F(1,19) = 34.659, p < 0.001, 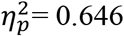). Responses in congruent trials were on average faster than in incongruent trials (mean RT difference: 105 ms). Critically, the amount of behavioral benefit depended on all other factors (4-way interaction: F(1.90,36.12) = 4.862, p = 0.015, 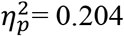). After model reduction, we found a significant interaction between *CONGRUENCE* and *STIMULATION* for ipsilateral gamma stimulation (F(1.93,36.65) = 4.578, 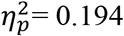). That means, responses to congruent stimuli were delayed by both in-and anti-phase ipsilateral gamma stimulation compared to sham (mean RT difference: 73 ms). In contrast, responses to incongruent stimuli did not differ between verum and sham (mean RT difference: 6 ms). This interaction was followed up by a non-parametric test based on CDFs where significance was determined by permutation statistics (corrected alpha: 0.000129, see *Methods* for details). Confirming ANOVA, differences between in-/anti-phase stimulation and sham were significant indicating reduced congruence effect due to stimulation (Fig. 3D; sham-in: 440-960 ms and 1330-1660 ms, sham-anti: 880-1610 ms). Additionally, we found significant differences between in-and anti-phase stimulation indicating increased reduction of congruence effect for early responses under in-phase stimulation (430-710 ms). Reproducing ANOVA patterns, CDFs of incongruent trials did not show significant differences between verum and sham. ANOVA also detected a significant interaction between *FREQUENCY* and *STIMULATION* for contralateral stimulation (F(1.78,33.83) = 3.771, p = 0.038, 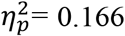). On average, differences between in-/anti-phase stimulation and sham were positive for alpha tACS (mean RT difference: 48 ms) and negative for gamma tACS (mean RT difference: −50 ms). In the post-hoc analysis, these differences were significant showing similar profiles for in-phase and anti-phase stimulation (Fig. 3E; Alpha in: 180-670ms and 1330-2280ms; Alpha anti: 450-690ms and 1010-2215ms; Gamma in: 525-970ms and 1330-2240ms; Gamma anti: 865-1015ms and 1615-2040 ms;). That is, there were no significant differences between in-and anti-phase stimulation.

Most participants reported tACS-related side effects (Fig. 3-2). While most participants reported “light” to “strong” skin sensations (median±inter quartile range; 1±1.25), only three participants reported phosphenes (0±0). Fatigue (0±1) and pain (0±1) were absent in the majority of participants. Importantly, the intensity of sensations overall did not differ with respect to sham, in-or anti-phase stimulation (uncorrected, for all, p > 0.09) and also showed no differences with respect to stimulation frequency (uncorrected, for all, p > 0.38). Next to the intensity of sensations, we asked for the time-course of a given sensation and coded responses into a binary decision for initial (0) or constant (1) stimulation (averages: 10 Hz: sham = .32, in = .68, anti = .74; 40 Hz: sham = .32, in = .68, anti = .63). Differences in the time-course of perception indicative of sham and verum condition were found significant or trending for both 10 and 40 Hz (all uncorrected; 10 Hz: sham-in: p = 0.03, sham-anti: p = 0.02, in-anti: p = 0.87; 40 Hz: sham-in: p = 0.03, sham-anti: p = 0.07, in-anti: p = 0.87). Sorensen-Dice similarity coefficients showed that ratings for 10 and 40 Hz stimulation were comparable (sham: 1, in: .77, anti: .77). Finally, correlations between the average effect of all three significant interactions detected with ANOVA and the overall intensity of skin sensations were weak and non-significant (all |r| < 0.25 and all p > 0.3). Interestingly, extreme values for skin sensations tended to occur more likely for participants with weak behavioral effects. Non-significant correlations thus showed opposite sign than would have been expected if the strength of sensations was indicative of behavioral effect size.

## Discussion

We investigated the putative role of long-range gamma synchronization between sensory cortices in multisensory perception. Importantly, spectral power in these regions was not modulated by the task. Instead, a previous whole-brain analysis of this EEG data showed that the power of alpha oscillations in frontal and parietal cortex was modulated by cross-modal attention (Anonymous, 2018). In the re-analysis of these data, we observed coherence between sensory gamma oscillations to be associated with cross-modal congruence. Furthermore, we tested the behavioral relevance of this sensory coupling by intervention with non-invasive brain stimulation. As hypothesized, behavioral benefits of cross-modal congruence were modulated by bifocal gamma hd-tACS when administered ipsilaterally to the hemisphere receiving sensory stimulation. Contralateral stimulation produced global changes in performance that showed opposite directions of behavioral effects for alpha and gamma stimulation.

### Cross-modal matching involves communication, not binding between modalities

In our paradigm, cross-modal matching between congruent inputs was associated with speeded responses when compared to the matching of incongruent inputs. This behavioral benefit of cross-modal congruence is well in line with previous studies that consistently showed faster responses and elevated accuracy of detecting or discriminating congruent multisensory stimuli (e.g. Bolognini et al., 2004; Schneider et al., 2008b; Göschl et al., 2014; Misselhorn et al., 2016). It was proposed that these behavioral benefits might arise because of cross-modal binding mediated by synchrony between sensory gamma oscillations (Senkowski et al., 2008). Accordingly, congruent multisensory stimuli should induce stronger cross-modal coupling when compared with incongruent stimuli. In our EEG data, we observed the opposite pattern of functional connectivity. That is, coherence of gamma oscillations in visual-tactile, audio-visual and audio-tactile networks was significantly reduced when participants were presented with congruent compared to incongruent stimuli. No differences in functional coupling were observed in the alpha band. Interestingly, this pattern of functional gamma coupling did not differ between attentional conditions. Thus, modulations of gamma band coherence were observed in all multisensory networks, irrespective of whether a given network was task-relevant or not. This finding indicates that the observed effects are predominantly stimulus-driven and less prone to top-down modulation. It should be noted, however, that the overall task-irrelevant audio-tactile network showed the tendency to be less strongly modulated by cross-modal congruence. We conclude that functional coupling during cross-modal matching, as implied in our paradigm, likely does not serve for cross-modal binding. Instead, the results can be interpreted in the context of the communication through coherence theory (Fries, 2009, 2015). In that perspective, coherence between gamma oscillations enables communication between interconnected cortical areas which is likely relevant for cross-modal matching. We speculate that that the increased need for communication between sensory areas in the case of incongruent stimuli was reflected by increased levels of cross-modal gamma coherence. Processing congruent stimuli might entail fast and efficient communication, and relatedly fast sensory decoupling that is reflected in decreased gamma coherence.

### Behavioral benefits of cross-modal congruence may involve fast sensory decoupling

In order to test the behavioral relevance of coupled sensory gamma oscillations for cross-modal matching, we analyzed behavioral changes induced with bifocal hd-tACS. As hypothesized, ipsilateral 40 Hz tACS reduced the behavioral benefit of cross-modal congruence, whereas 10 Hz stimulation did not show significant effects. This reduction of the congruence-effect during 40 Hz tACS was due to significantly increased response latencies for congruent trials, while incongruent trials showed no difference between verum and sham. Finding incongruent trials unchanged by tACS, might suggest that the strength of communication cannot be improved by tACS. That is, as long as effective communication between sensory cortices is established, coherence is near an optimal point and does not benefit from external synchronization. As proposed above, it is possible that the behavioral benefit of processing congruent stimuli partly relies on fast decoupling and thus efficient further processing in regions of multisensory convergence such as the temporal or parietal cortex. Finding responses to congruent trials delayed by gamma tACS is consistent with this idea. Accordingly, sensory decoupling might have been hampered by gamma tACS which may have kept channels for communication open. Collectively, these findings lend additional support to the idea that gamma oscillations play a critical role in structuring interactions and likely communication between sensory cortical areas.

Surprisingly, both in-and anti-phase stimulation showed on average similar effects. As suggested by previous work, we expected that in-phase stimulation would synchronize cortical activity under the bifocal montage while anti-phase stimulation would desynchronize activity (Polanía et al., 2012; Helfrich et al., 2014; Schwab et al., 2018). In contrast to these studies, we did not aim to modulate coupling between association cortices, but directly between sensory regions of distinct modalities. It is possible that – unlike communication within sensory modalities – multisensory communication is rather broadly tuned in frequency. In our EEG data, gamma frequencies of maximal coupling varied between 60 and 90 Hz without showing a clear peak on group level (mean peak frequency: 77.19 Hz, see also Fig. 2-1). This is consistent with the literature where gamma peak frequency usually shows large inter-individual variability (e.g. Schwarzkopf et al., 2012). Moreover, bandwidth of sensory gamma oscillations varies between modalities in a range between 30 and more than 100 Hz (*visual:* Jia et al. 2011; Friese et al. 2016; Sumner et al. 2018; *somatosensory:* Wahnoun et al. 2015; Ryun et al. 2017; von Lautz et al. 2017; *auditory:* Edwards et al. 2005; Griffiths et al. 2010; Mulert et al. 2011). Given this large variability within and across modalities, narrow tuning of direct multisensory communication channels would, in fact, be surprising. It is thus possible that cross-modal interactions are broadly tuned, and that 40 Hz tACS in this study produced rather broad-band than frequency-specific effects.

Alternatively, tACS could have produced local effects in gamma power that did not differ between in-and anti-phase stimulation. While we cannot exclude this possibility, differential modulations on the level of response time distributions argue against this explanation: while in-phase stimulation predominantly affected early responses (~500-1000 ms), anti-phase stimulation affected later responses (~900-1600 ms). This led to a significantly stronger negative influence of in-phase compared with anti-phase stimulation for responses between 430 and 710 ms. While this is in line with our interpretation of prevented fast decoupling, largely comparable CDF differences point out that our initial hypothesis about inverse effects of in-and anti-phase stimulation is clearly not supported by the present results. This hypothesis assumed that zero-phase lag coherence should be beneficial for communication between modalities. Considering conduction delays between sensory cortices and peak frequency differences, optimal phase angles for cross-modal communication need not necessarily be zero and might show large variability.

### Contralateral alpha/gamma dynamics regulate cortical excitability

In order to control for unspecific effects of tACS, we used contralateral stimulation as an active control. Contrary to our expectations, these modulations of task-irrelevant cortical regions showed global modulations of behavior that were specific with respect to frequency. While 40 Hz stimulation showed overall slowing of responses, 10 Hz stimulation showed overall speeding of responses. These findings can be related to the functionally opposing roles of alpha and gamma dynamics in cortex: while gamma oscillations are enhanced from activated cortical areas (Donner and Siegel, 2011), alpha oscillations predominate in task-irrelevant cortical regions (Jensen and Mazaheri, 2010). This view is supported by a negative respectively positive correlation of EEG gamma and alpha power with the BOLD signal (Mulert et al., 2010; Scheeringa et al., 2011). Importantly, the level of ongoing alpha activity could be shown to be a readout of cortical excitability as determined by transcranial magnetic stimulation (Romei et al., 2008). Cortical excitability, as controlled by alpha/gamma dynamics, is also discussed as a mechanism underlying top-down control of perceptual processes (Jensen and Mazaheri, 2010; Bonnefond and Jensen, 2015). For instance, cued spatial attention led to a lateralization of pre-stimulus alpha power to the task-irrelevant hemisphere, while stimulus-related gamma activity was lateralized to the task-relevant hemisphere (Marshall et al., 2015). In our tACS experiment, stimuli were presented lateralized and pre-trial cues were used to guide spatial attention of participants. We propose that reducing cortical excitability of the task-irrelevant hemisphere through alpha stimulation improved processing in the task-relevant hemisphere. Conversely, gamma stimulation might have increased excitability of the task-irrelevant hemisphere and thereby disrupted processing in the task-relevant hemisphere. A similar result has recently been obtained for unilateral stimulation over temporo-parietal cortex in a dichotic listening task (Wöstmann et al., 2018). In their study, alpha stimulation decreased recall of contralateral items while gamma stimulation showed the opposite effect. Our results, thus, add to the abundant literature that suggests an important role of alpha/gamma dynamics in modulating cortical excitability.

### Limitations and future directions

Despite the complementary nature of the two presented experiments, investigating two distinct groups of participants was a limiting factor. Thus, we were not able to tailor the tACS stimulation frequencies individually based on preceding EEG recordings. However, using canonical stimulation frequencies, our data points out that such tailoring might not be necessary because tACS in the gamma range might have broad instead narrow-banded effects. Future studies addressing the spectral profile of tACS-related changes in cortical activity will be helpful. Furthermore, analyses aimed at finding correlations between tACS-modulated behavior and electrophysiology were not possible. Ideally, such relations should be established within the same dataset. Firm conclusions on how cross-modal interactions relate to behavior as well as how (high-frequency) tACS influences neurophysiology depend on such data. Analyzing acute effects of tACS, however, is hampered by the unsolved problem of correcting tACS-related artifacts in electrophysiological recordings (Noury et al., 2016). Meanwhile, future studies could resort to intermittent stimulation protocols that make use of potentially occurring aftereffects.

## Conclusions

Our EEG results provide evidence that coordinated sensory gamma oscillations play an important role for direct cross-modal interactions. We suggest that, much like interactions within sensory streams, phase-coupled gamma oscillations provide the functional scaffold for cross-modal communication. Findings from the tACS experiment corroborate this notion by showing frequency-specific effects on behavioral responses for congruent stimuli. The pattern of results indicates that efficient matching between modalities might involve flexible coupling and decoupling of gamma oscillations between sensory cortical areas. The absence of clear phase-specific effects of this stimulation regime, however, suggests that both acute effects of tACS and cross-modal interactions are poorly understood. Finally, we provide evidence for the idea that lateralized alpha/gamma dynamics are related to fluctuations of cortical excitability underlying selective attention.

## Acknowledgements

This work was supported by grants from the DFG (SFB 936/A3/Z1, SFB TRR 169/B1 and SPP 1665/EN 533/13-1 to AKE; SPP 1665/SCHN 1511/1-2 to TRS) and the EU (ERC-2010-AdG-269716 to AKE). The authors would like to thank Karin Deazle for help with recruitment of participants and data recording.

## Extended data

**Figure 2-1.**
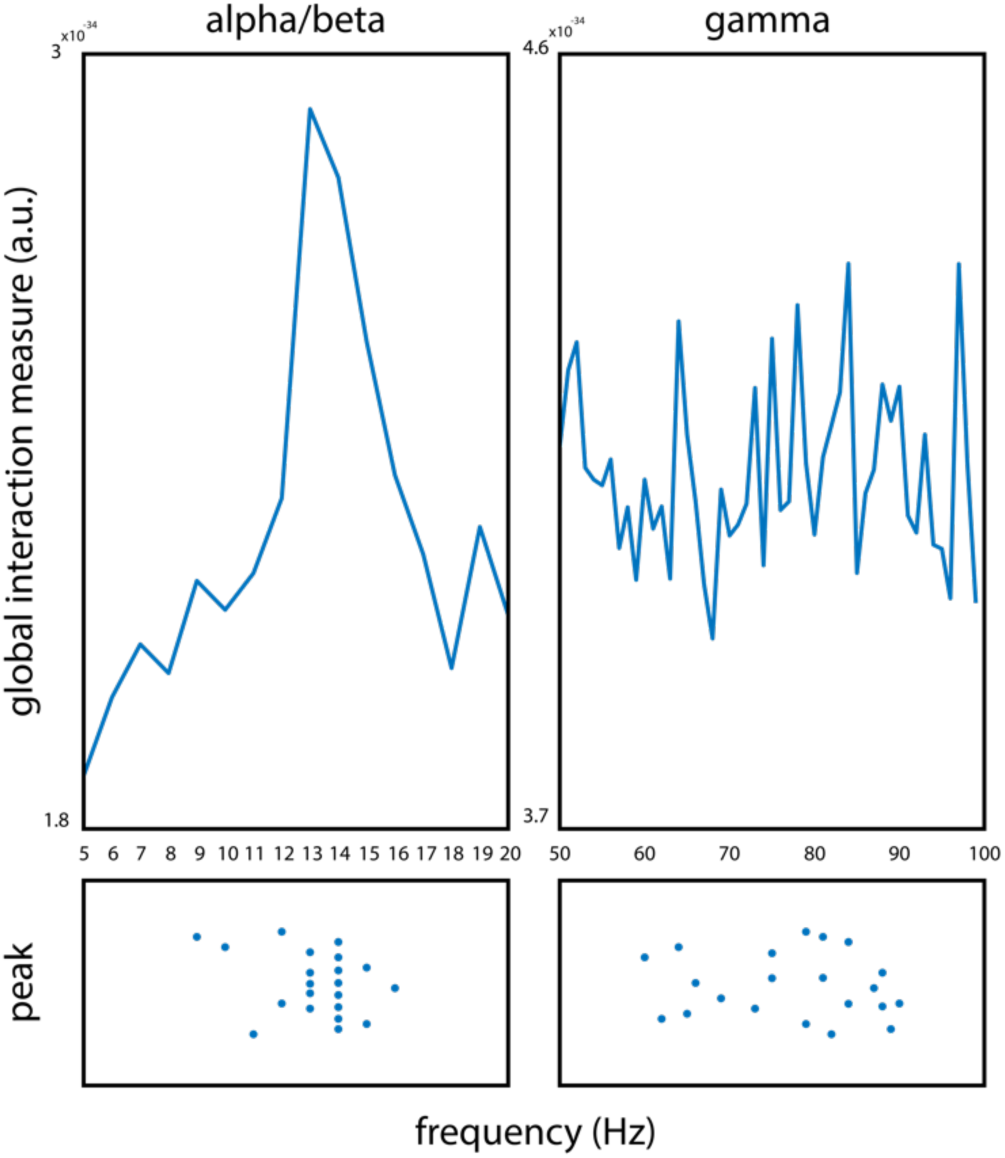
Global interaction measure. **Spectra and individual peaks of GIM for sensory space EEG data.** (top left) Spectrum from 5 to 20 Hz showing a clear group level peak at 13 Hz. (bottom left) Distribution of individual peak frequencies within a band from 8 to 20 Hz. (top right) Spectrum from 50 to 100 Hz does not show a distinct group level peak. (bottom right) Distribution of individual peak frequencies within a band from 60 to 90 Hz.

**Figure 2-2.**
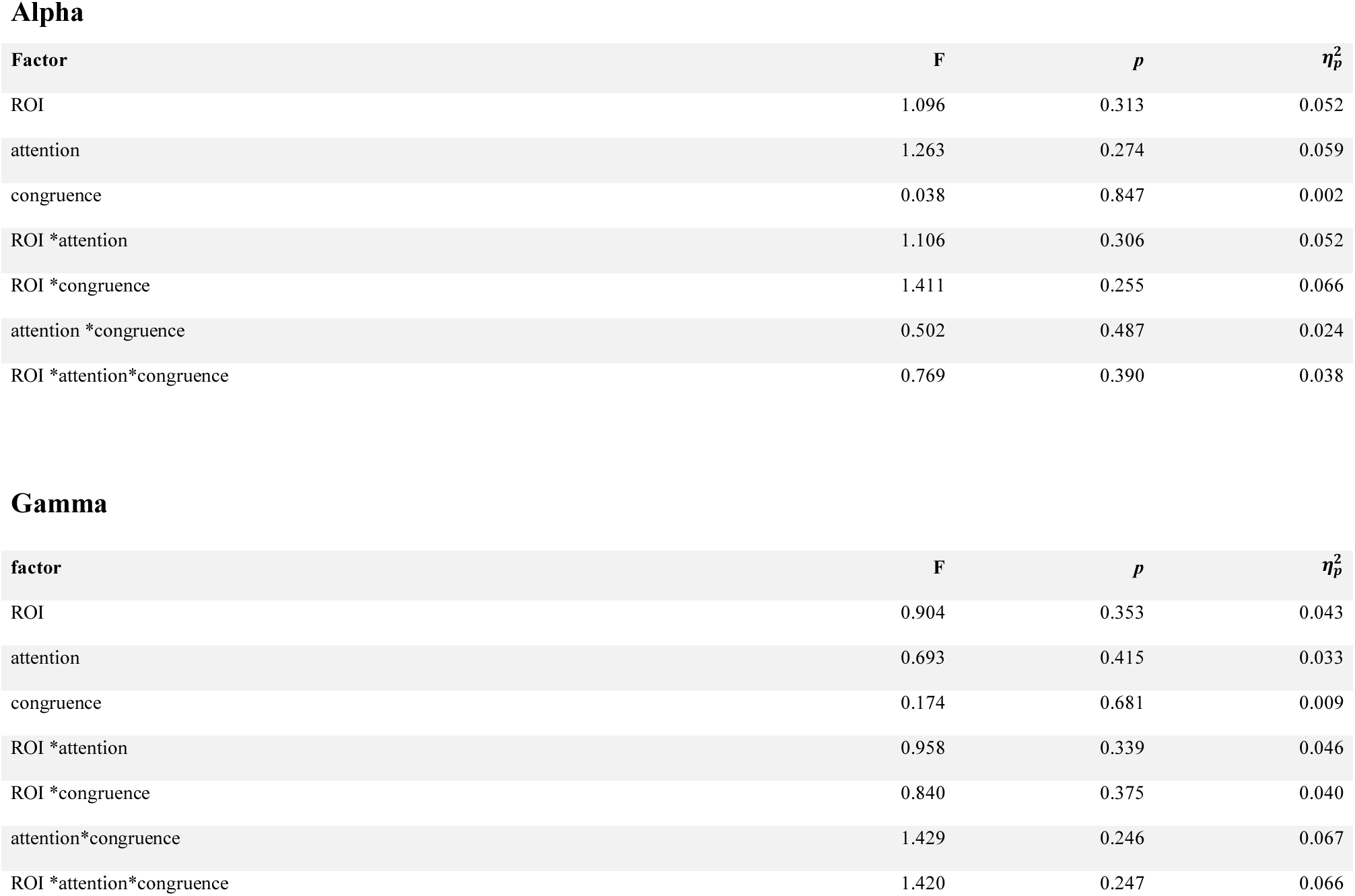
Complete results from ANOVA on spectral power.

**Figure 2-3.**
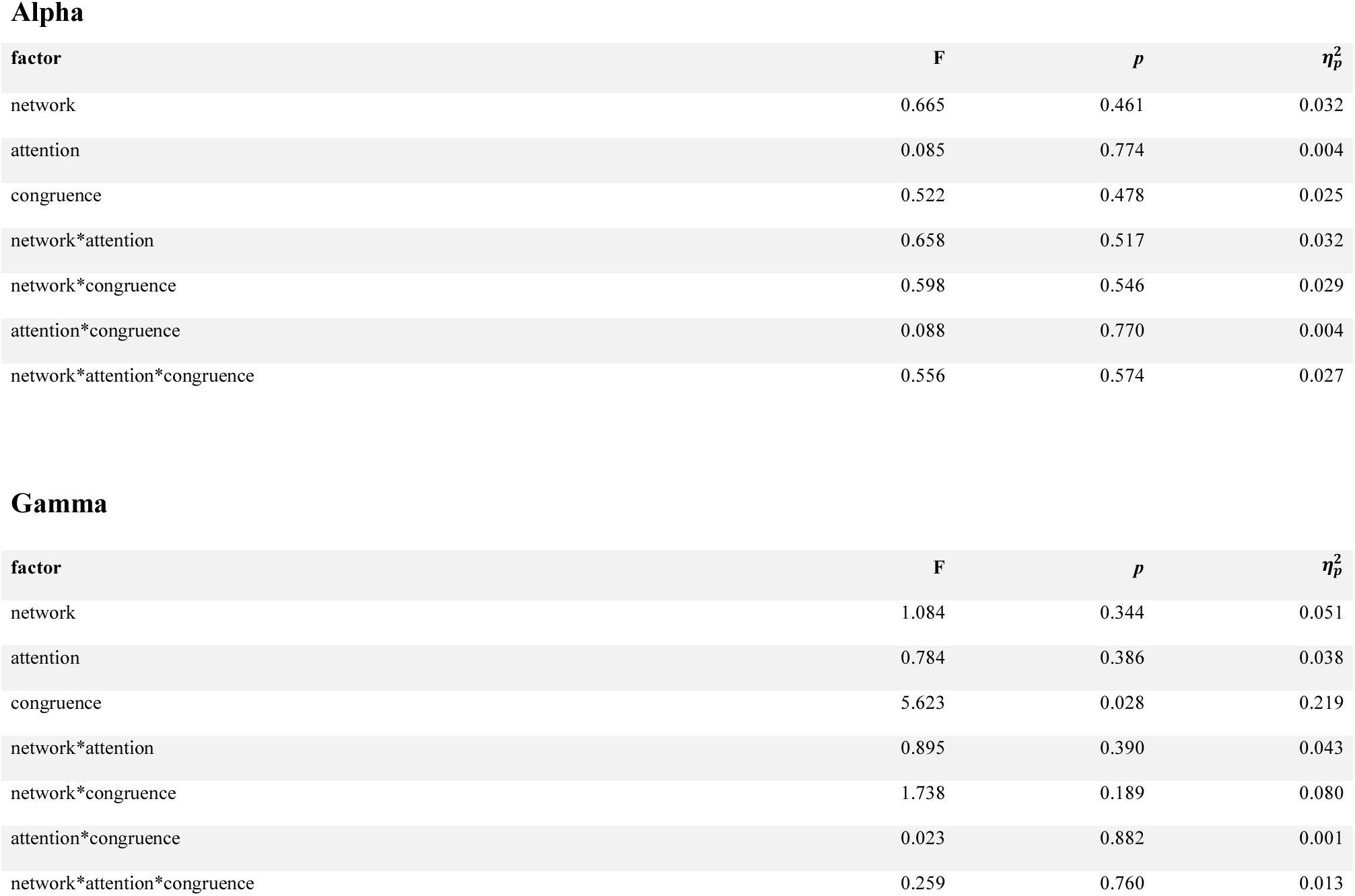
Complete results from ANOVA on phase coherence.

**Figure 3-1.**
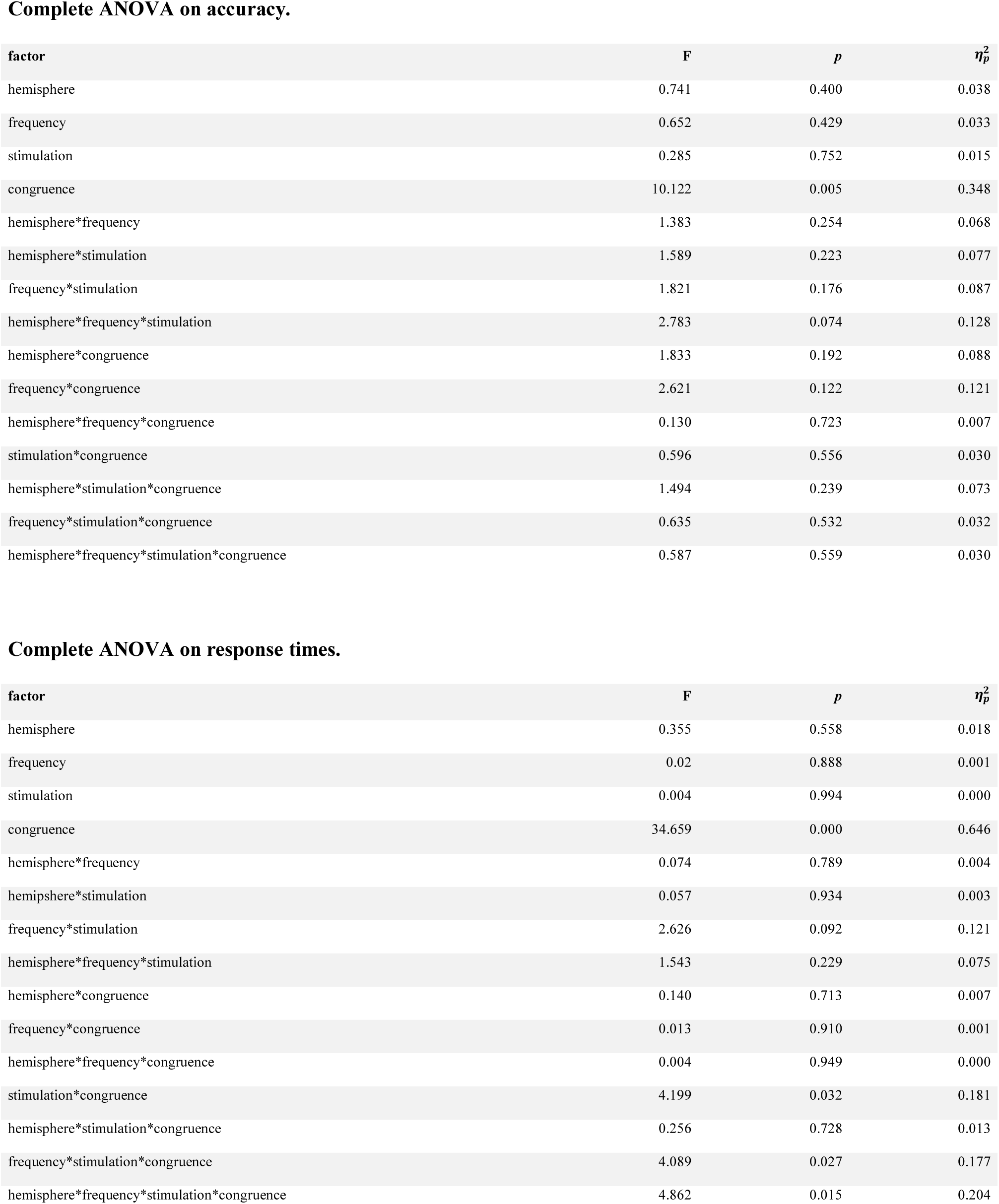
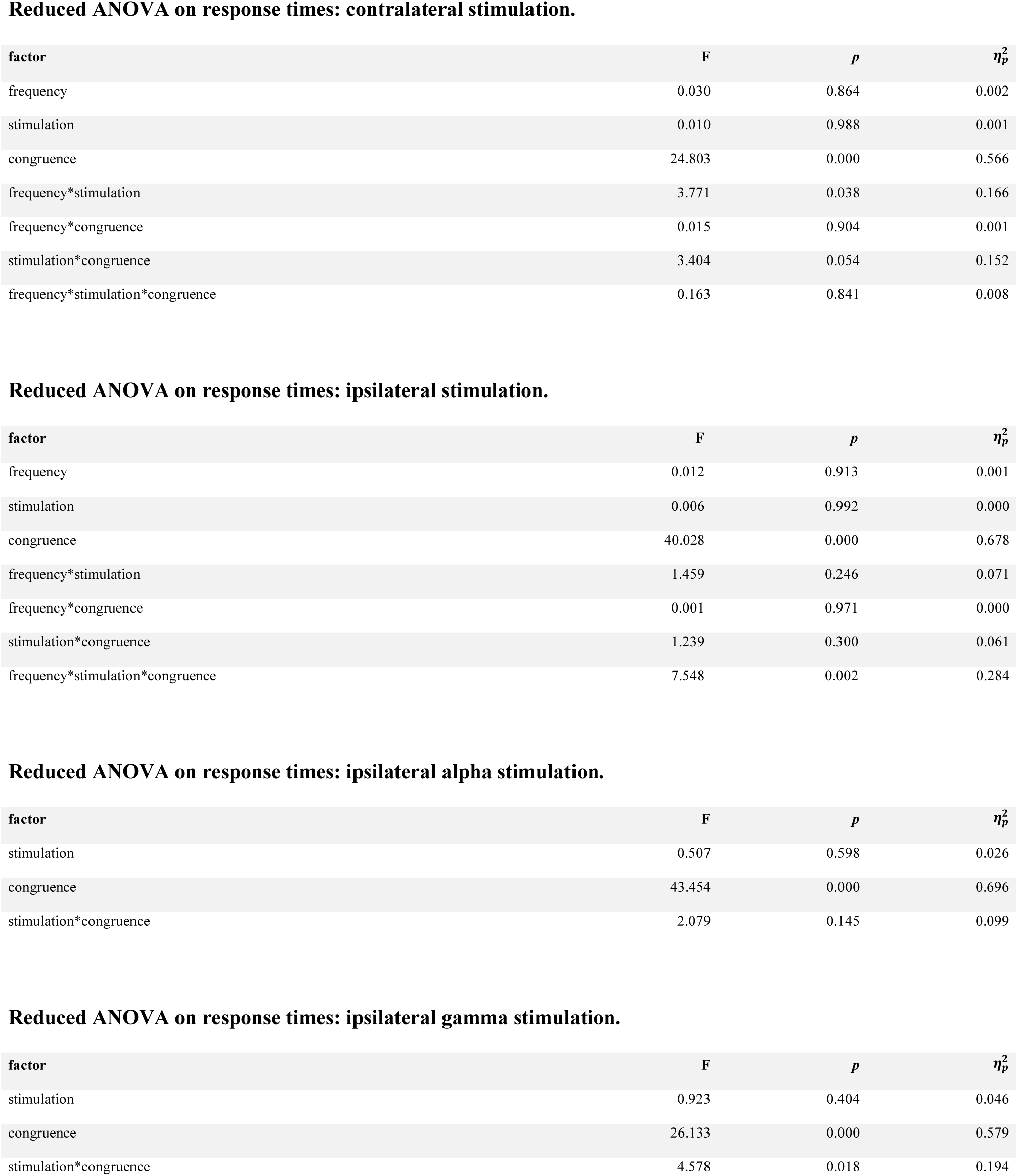
Complete and reduced ANOVA results from behavior in tACS experiment.

**Figure 3-2.**
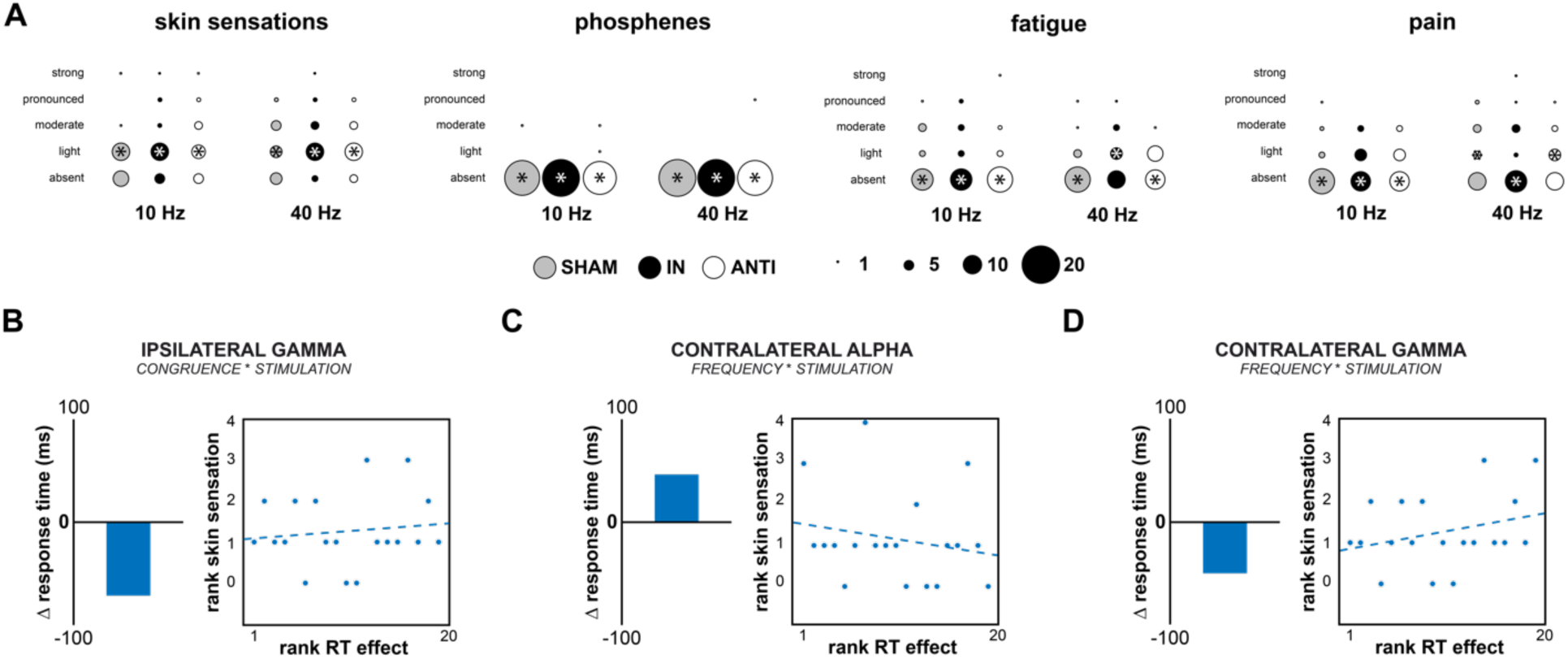
Side effects of tACS. **(A)** Visualization of questionnaire data for skin sensations (aggregated across itching, warmth, stinging, pulsating), phosphenes, fatigue and pain. Lowest row represents “absent” response while upper rows indicate “light” to “strong” sensation. Size of circles represent number of responses and asterisks indicate median response per condition and sensation. **(B-D)** Correlations of ranked behavioral effect detected in ANOVA with skin sensations (rank 1 is lowest value). Bar plots indicate direction of behavioral effects. Across all effects, correlations are weak and non-significant, but show signs that are opposite to what would have been expected if side effects drove the behavioral effects.

## References

Balz J, Romero YR, Keil J, Krebber M, Niedeggen M, Gallinat J, Senkowskil D (2016) Beta/Gamma Oscillations and Event-Related Potentials Indicate Aberrant Multisensory Processing in Schizophrenia. Front Psychol 7:1896.

Bastos AM, Schoffelen JM (2016) A Tutorial Review of Functional Connectivity Analysis Methods and Their Interpretational Pitfalls. Front Syst Neurosci 9:175.

Bolognini N, Frassinetti F, Serino A, Làdavas E (2004) “Acoustical vision” of below threshold stimuli: interaction among spatially converging audiovisual inputs. Exp Brain Res 160:273–282.

Bonnefond M, Jensen O (2015) Gamma Activity Coupled to Alpha Phase as a Mechanism for Top-Down Controlled Gating. PLoS ONE 10:e0128667.

Botev ZI, Grotowski JF, Kroese DP (2010) Kernel density estimation via diffusion. Ann Stat 38:2916–2957.

Curic S, Leicht G, Thiebes S, Andreou C, Polomac N, Eichler IC, Eichler L, Zöllner C, Gallinat J, Steinmann S, Mulert C (2019) Reduced auditory evoked gamma-band response and schizophrenia-like clinical symptoms under subanesthetic ketamine. Neuropsychopharmacology.

Desikan RS, Ségonne F, Fischl B, Quinn BT, Dickerson BC, Blacker D, Buckner RL, Dale AM, Maguire RP, Hyman BT, Albert MS, Killiany RJ (2006) An automated labeling system for subdividing the human cerebral cortex on MRI scans into gyral based regions of interest. NeuroImage 31:968–980.

Donner TH, Siegel M (2011) A framework for local cortical oscillation patterns. Trends Cogn Sci 15:191–199.

Edwards E, Soltani M, Deouell LY, Berger MS, Knight RT (2005) High gamma activity in response to deviant auditory stimuli recorded directly from human cortex. J Neurophysiol 94:4269–4280.

Engel AK, Konig P, Kreiter AK, Singer W (1991) Interhemispheric synchronization of oscillatory neuronal responses in cat visual cortex. Science 252:1177–1179.

Engel AK, Fries P, Singer W (2001) Dynamic predictions: oscillations and synchrony in top-down processing. Nat Rev Neurosci 2:704–716.

Ewald A, Marzetti L, Zappasodi F, Meinecke FC, Nolte G (2012) Estimating true brain connectivity from EEG/MEG data invariant to linear and static transformations in sensor space. NeuroImage 60:476–488.

Fries P (2009) Neuronal gamma-band synchronization as a fundamental process in cortical computation. Annu Rev Neurosci 32:209–224.

Fries P (2015) Rhythms for Cognition: Communication through Coherence. Neuron 88:220–235.

Friese U, Daume J, Göschl F, König P, Wang P, Engel AK (2016) Oscillatory brain activity during multisensory attention reflects activation, disinhibition, and cognitive control. Sci Rep 6:32775.

Ghazanfar AA, Schroeder CE (2006) Is neocortex essentially multisensory? Trends Cogn Sci 10:278–285.

Gray CM, König P, Engel AK, Singer W (1989) Oscillatory responses in cat visual cortex exhibit inter-columnar synchronization which reflects global stimulus properties. Nature 338:334–337.

Griffiths TD, Kumar S, Sedley W, Nourski KV, Kawasaki H, Oya H, Patterson RD, Brugge JF, Howard MA (2010) Direct recordings of pitch responses from human auditory cortex. Curr Biol CB 20:1128–1132.

Goldman RI, Stern JM, Engel J, Cohen MS (2002) Simultaneous EEG and fMRI of the alpha rhythm. Neuroreport 13:2487–2492.

Göschl F, Engel AK, Friese U (2014) Attention modulates visual-tactile interaction in spatial pattern matching. PloS One 9:e106896.

Helfrich RF, Knepper H, Nolte G, Strüber D, Rach S, Herrmann CS, Schneider TR, Engel AK (2014) Selective Modulation of Interhemispheric Functional Connectivity by HD-tACS Shapes Perception. PLoS Biol 12:e1002031.

Jensen O, Mazaheri A (2010) Shaping Functional Architecture by Oscillatory Alpha Activity: Gating by Inhibition. Front Hum Neurosci 4:186.

Jia X, Smith MA, Kohn A (2011) Stimulus selectivity and spatial coherence of gamma components of the local field potential. J Neurosci Off J Soc Neurosci 31:9390–9403.

Krebber M, Harwood J, Spitzer B, Keil J, Senkowski D (2015) Visuotactile motion congruence enhances gamma-band activity in visual and somatosensory cortices. NeuroImage 117:160–169.

Marshall TR, O’Shea J, Jensen O, Bergmann TO (2015) Frontal eye fields control attentional modulation of alpha and gamma oscillations in contralateral occipitoparietal cortex. J Neurosci Off J Soc Neurosci 35:1638–1647.

Misselhorn J, Daume J, Engel AK, Friese U (2016) A matter of attention: Crossmodal congruence enhances and impairs performance in a novel trimodal matching paradigm. Neuropsychologia 88:113–122.

Mulert C, Leicht G, Hepp P, Kirsch V, Karch S, Pogarell O, Reiser M, Hegerl U, Jager L, Moller HJ, McCarley RW (2010): Single-trial coupling of the gamma-band response and the corresponding BOLD signal. Neuroimage 49:2238–2247.

Mulert C, Kirsch V, Pascual-Marqui R, McCarley RW, Spencer KM (2011) Long-range synchrony of γ oscillations and auditory hallucination symptoms in schizophrenia. Int J Psychophysiol Off J Int Organ Psychophysiol 79:55–63.

Nolte G, Bai O, Wheaton L, Mari Z, Vorbach S, Hallett M (2004) Identifying true brain interaction from EEG data using the imaginary part of coherency. Clin Neurophysiol Off J Int Fed Clin Neurophysiol 115:2292–2307.

Nolte G, Dassios G (2005) Analytic expansion of the EEG lead field for realistic volume conductors. Phys Med Biol 50:3807–3823.

Noury N, Hipp JF, Siegel M (2016) Physiological processes non-linearly affect electrophysiological recordings during transcranial electric stimulation. Neuroimage 140:99–109.

Pascual-Marqui RD, Lehmann D, Koukkou M, Kochi K, Anderer P, Saletu B, Tanaka H, Hirata K, John ER, Prichep L, Biscay-Lirio R, Kinoshita T (2011) Assessing interactions in the brain with exact low-resolution electromagnetic tomography. Philos Transact A Math Phys Eng Sci 369:3768–3784.

Patel J, Bansal V, Minha P, Ho J, Datta A, Bikson M (2009) High-Density Transcranial Direct Current Stimulation (HD-tDCS): Skin Safety and Comfort. J Med Devices 3:027554.

Polanía R, Nitsche MA, Korman C, Batsikadze G, Paulus W (2012) The Importance of Timing in Segregated Theta Phase-Coupling for Cognitive Performance. Curr Biol 22:1314–1318.

Romei V, Brodbeck V, Michel C, Amedi A, Pascual-Leone A, Thut G (2008) Spontaneous Fluctuations in Posterior α-Band EEG Activity Reflect Variability in Excitability of Human Visual Areas. Cereb Cortex 18:2010–2018.

Ryun S, Kim JS, Lee H, Chung CK (2017) Tactile Frequency-Specific High-Gamma Activities in Human Primary and Secondary Somatosensory Cortices. Sci Rep 7:15442.

Saturnino GB, Antunes A, Thielscher A (2015) On the importance of electrode parameters for shaping electric field patterns generated by tDCS. NeuroImage 120:25–35.

Schwarzkopf DS, Robertson DJ, Song C, Barnes GR, Rees G (2012) The frequency of visually induced gamma-band oscillations depends on the size of early human visual cortex. J Neurosci Off J Soc Neurosci 32:1507–1512.

Spence C (2011) Crossmodal correspondences: A tutorial review. Atten Percept Psychophys 73:971–995.

Scheeringa R, Fries P, Petersson KM, Oostenveld R, Grothe I, Norris DG, Hagoort P, Bastiaansen MC (2011): Neuronal dynamics underlying high-and low-frequency EEG oscillations contribute independently to the human BOLD signal. Neuron 69:572–583.

Schneider TR, Debener S, Oostenveld R, Engel AK (2008a) Enhanced EEG gamma-band activity reflects multisensory semantic matching in visual-to-auditory object priming. NeuroImage 42:1244–1254.

Schneider TR, Engel AK, Debener S (2008b) Multisensory Identification of Natural Objects in a Two-Way Crossmodal Priming Paradigm. Exp Psychol 55:121–132.

Schwab BC, Misselhorn J, Engel AK (2018) Modulation of large-scale cortical coupling by transcranial alternating current stimulation. bioRxiv:484014.

Senkowski D, Talsma D, Herrmann CS, Woldorff MG (2005) Multisensory processing and oscillatory gamma responses: effects of spatial selective attention. Exp Brain Res 166:411–426.

Senkowski D, Talsma D, Grigutsch M, Herrmann CS, Woldorff MG (2007) Good times for multisensory integration: Effects of the precision of temporal synchrony as revealed by gamma-band oscillations. Neuropsychologia 45:561–71.

Senkowski D, Schneider TR, Foxe J, Engel AK (2008) Crossmodal binding through neural coherence: implications for multisensory processing. Trends Neurosci 31:301–409.

Sumner RL, McMillan RL, Shaw AD, Singh KD, Sundram F, Muthukumaraswamy SD (2018) Peak visual gamma frequency is modified across the healthy menstrual cycle. Hum Brain Mapp 39:3187–3202.

Stone DB, Coffman BA, Bustillo JR, Aine CJ, Stephen JM (2014) Multisensory stimuli elicit altered oscillatory brain responses at gamma frequencies in patients with schizophrenia. Front Hum Neurosci 8:788.

Tallon-Baudry C, Bertrand O, Delpuech C, Pernier J (1996) Stimulus Specificity of Phase-Locked and Non-Phase-Locked 40 Hz Visual Responses in Human. J Neurosci 16:4240–4249.

Treisman A (1996) The binding problem. Curr Opin Neurobiol 6:171–178.

Uhlhaas PJ, Singer W. (2010) Abnormal neural oscillations and synchrony in schizophrenia. Nat Rev Neurosci 11:100–113.

von Lautz AH, Herding J, Ludwig S, Nierhaus T, Maess B, Villringer A, Blankenburg F (2017) Gamma and Beta Oscillations in Human MEG Encode the Contents of Vibrotactile Working Memory. Front Hum Neurosci 11:576.

Watson AB, Pelli DG (1983) QUEST: a Bayesian adaptive psychometric method. Percept Psychophys 33:113–120.

Wahnoun R, Benson M, Helms-Tillery S, Adelson PD (2015) Delineation of somatosensory finger areas using vibrotactile stimulation, an ECoG study. Brain Behav 5:e00369.

Wöstmann M, Vosskuhl J, Obleser J, Herrmann CS (2018) Opposite effects of lateralised transcranial alpha versus gamma stimulation on auditory spatial attention. Brain Stimulat 11:752–758.

Yuval-Greenberg S, Deouell LY (2007) What you see is not (always) what you hear: induced gamma band responses reflect cross-modal interactions in familiar object recognition. J Neurosci 27:1090–1096.

